# GVC: An ultra-fast and all-round genome variant caller

**DOI:** 10.1101/182089

**Authors:** Zhongbo Zhang, Longhui Yin, Lingtong Hao, Lihua Cao, Changlin Liu, Mengzhu Chen, Cheng Li, Zhao Xu, Shaoping Ling

## Abstract

Genome variant detection is a challenge task in cancer genome analysis. Current software for variant detection is very time-consuming and in the same time not accurate enough to satisfy the requirements of clinical applications in precision oncology. We developed an all-round ultra-fast Genomic Variant Caller (GVC), which can detect Germline and Somatic variants, including SNVs and INDELs from genome sequencing data with super high speed and accuracy. GVC achieved mean F1-measure of 99.34% and 97.92% for Germline SNVs in WGS and WES datasets, respectively. And GVC reached mean F1-measure of 97.34% and 89.03% for Germline INDELs in WGS and WES datasets, respectively. GVC achieved mean F1-measure of 90.79% and 86.3% for Somatic SNVs in WGS and WES datasets, respectively. GVC reached mean F1-measure of 76.88% and 68.36% for Somatic INDELs in WGS and WES datasets, respectively. GVC outperformed all the other widely used pipelines. GVC showed super-fast processing speed, exhibiting 128-fold and 54-fold increase in comparison to GATK in WGS and WES datasets processing, respectively. GVC webtool is available from GVC Web Tool https://gvc.0cancer.cn. Basing on the accurate GVC algorithm, we also released tumor mutation burden (TMB) results of TCGA datasets on GVC Web Tool.

## INTRODUCTION

Detection of the Germline and Somatic mutation is key step in characterizing the blood and tissue genome. Germline mutations are those DNA sequence changes occur during various stages of zygote development exist in all kinds of tissues that can be passed on to offspring. Germline mutations occur in tumor suppressor genes or proto-oncogenes predispose an individual to develop tumor. Identification of critical cancer-gene variants in tumor samples could better defines patient diagnosis and prognosis, and it also presents what targeted therapies must be administered to improve the care of selected cancer patients in the personalized-medicine scenario. The expanded applications of NGS in oncology include not only prediction of checkpoint inhibitor response, quantification of tumor mutational burden and neoantigen burden, but also the personalization of cell transfer technologies and cancer vaccines^1^.

Next generation sequencing (NGS) is by far the most promising technology for de novo mutation detection. However two primary obstacles hinder the clinical application of mutation detection via NGS. One is the precision of variant calling with NGS data. Mutations are difficult to identify accurately due to low occurring frequency, tumor purity and clonality^2^. The PCAWG (Pan-Cancer Analysis of Whole Genomes) project is an international collaboration from ICGC (the International Cancer Genome Consortium), investigating patterns of mutation in more than 2,800 patients with cancer. INDELs results in PCAWG project called by extensively used bioinformatic pipelines from Sanger, DKFZ/EMBL and Broad institute represent a low level of consistency at 42%. In order to achieve a higher variant call accuracy, many algorithm strategies have been designed and implemented, including the heuristic approaches (VarScan2, Shimmer, and RADIA etc)^3-5^, joint genotype analysis (SomaticSniper, FaSDsomatic, and SAMtools etc)^6,7^, joint allele frequencies analysis (Strelka, MuTect, and LoFreq etc)^8-10^, haplotype-based strategy (Platypus, FreeBayes, and MuTect2 etc)^4^, and machine learning methods (MutationSeq, SomaticSeq, and SNooPer etc)^11-13^. Variant calling using machine learning methods is essentially a classification problem. Four train classifiers including random forest, Bayesian adaptive regression tree, support vector machine, and logistic regression have already been explored in variant detection. However, the overall performance is still not ideal.

The other major obstacle of NGS data processing in clinical application is variant calling speed. Technical advances in NGS technologies have reduced both the cost and time required to generate a large volume of sequencing data, but variant calling is still such a time-consuming process due to the high complexity of the issue and the large volume of the data. In mutation detection, widespread used software such as Bcftools^14^, Varscan^4^, and GATK^15^, all run at a very slow speed to detect variants, which can hardly satisfy the requirement of clinical applications. It is a great challenge to obtain variant calling results efficiently and accurately from massive data.

We developed GVC (genome variant caller), which can detect Germline and Somatic mutations with ultra-fast speed and high accuracy. The core algorithm of GVC is integrated with Feature Space construction and XGBoost machine learning. XGBboost is a scalable end-to-end tree boosting system, and it scales beyond billions of examples using far fewer resources than existing systems and exhibits tremendous potential in variant detection. We applied several benchmarking datasets including whole genome and whole exon sequencing datasets from ICGC and TCGA to evaluate the sensitivity and specificity of GVC in comparison to other widespread used softwares. Results validated that GVC could deliver genome variant caller with high accuracy ultra-fast speed.

## RESULTS AND DISCUSSION

### Overview of the GVC workflow

As a machine-learning based pipeline, GVC can be divided into two parts (Figure 1), Genome Analysis and Training. After cleaning of the raw sequencing data, alignment of the fastq-format reads to a reference genome (GRCh37) is carried out by BWA-mem to obtain corresponding BAM (Binary Alignment Map) files. GVC takes BAM files as input for either only normal sample or matched normal-tumor sample pairs. Features fitted to different applicable scenario are extracted automatically. False variants were filtered through machine-learning model to obtain the candidate mutations. In the training part of the GVC pipeline, uncertain variants are validated by Sanger sequencing or ultra-deep sequencing. Validated variants and datasets are added into the iGC database which can be used to train updated machine learning model for better performance. Detecting, validation and training compose the core of the GVC workflow, and make it a flexible, scalable and customizable.

**Figure 1.**
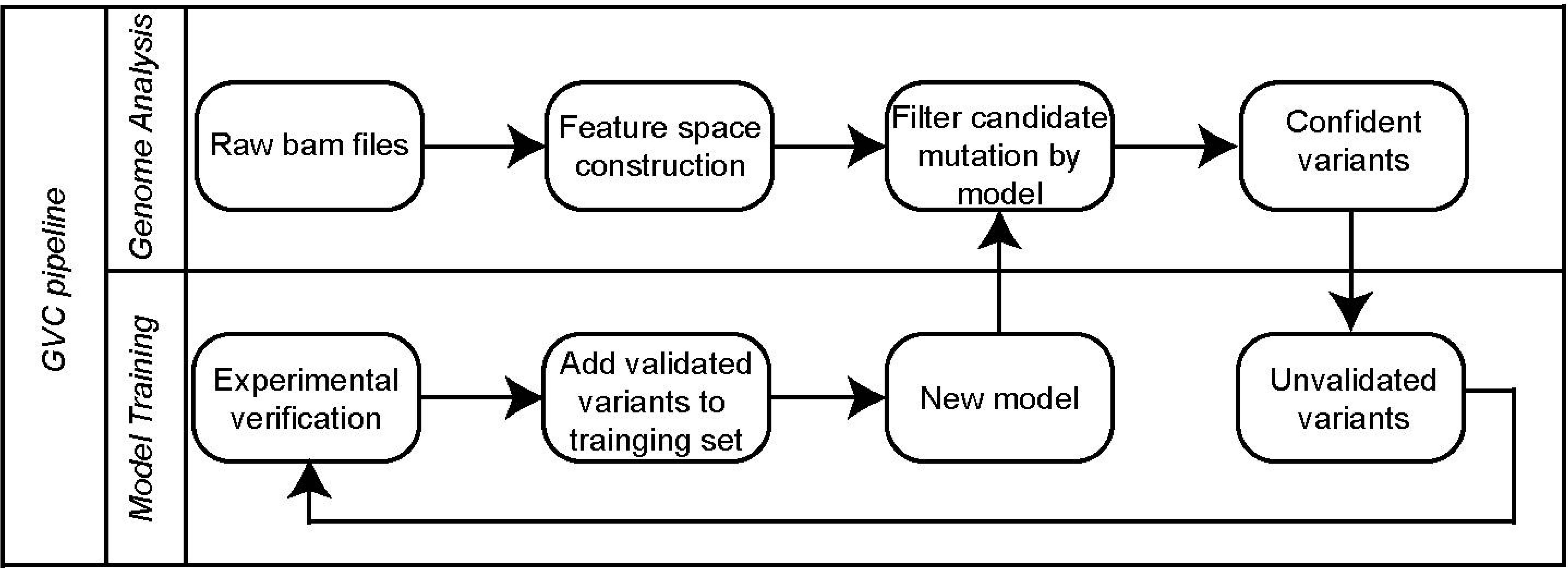
Overview of the GVC pipeline. GVC pipeline can be divided into two main procedures, (i) GVC takes sorted BAM as input files and constructs feature vectors in order to pick out confident variants, (ii) GVC elevates model accuracy by continuously adding validated data.

GVC exhibits two main characteristics. Firstly it applied BAM files to extracted features of sequencing data, such as read depth, mapping quality, base quality and so on (see method). Around 30 Features could be extracted for Somatic variant calling. It is notable that Feature Space construction of GVC is a flexible and scalable process according to diverse applied scenarios. For example in liquid biopsy sample, GVC added at least 10 features to Feature Space in order to eliminate recurrent artifacts and stochastic errors. Secondly GVC is a machine learning-based variant caller. Distinguished from other machine-learning based variant callers, GVC applied XGBoost classifier as its core algorithm. According to the Feature Space or experimental validation, GVC could adjust a set of “ground truth” variants, and retrained the model to get better performance.

It should be pointed out that various library construction methods, diverse sequencing platforms, different quality evaluation systems, sequencing depths, and read lengths, all contribute to diversity of data features. We suggested that users select appropriate GVC models or train their GVC models with their own data (see method) to achieve better performance. We believe GVC has broad application potential due to its scalable Feature Space and customizable data models.

### Accuracy of GVC on Germline variant calling

Two WGS (Whole Genome Sequencing) and four WES (Whole Exome Sequencing) training datasets derived from well-studied human sample NA12877 and NA12878 were downloaded from Platinum Genomes project. A reference paper^16^, and k-fold cross validation strategy was used in training process (see method and Supplementary Table 1). We assessed the accuracy of GVC’s SNVs and INDELs calls against the Genome in a Bottle Consortium (GIAB) truth set derived from NA12877 (3,520,925 SNVs and 512,092 INDELs) and NA12878 (3,526,588 SNVs and 522,924 INDELs). GVC achieved F1-measure of 99.34% and 97.92% for Germline SNVs in WGS and WES datasets, respectively. These detection rates exceeded those of Bcftools, GATK and Varscan tools (WGS: ≤99.19%, WES: ≤97.57%) (Figure 2A, Supplementary Table 2, Supplementary Table 3 and Supplementary Fig. 1). And GVC also reached F1-meature of 97.34% and 89.03% for Germline INDELs in WGS and WES datasets, respectively. These detection rates exceeded those of Bcftools, GATK and Varscan tools (WGS: ≤95.73%, WES: ≤78.96%) (Figure 2B, Supplementary Table 2, Supplementary Table 3 and Supplementary Fig. 1). In short, GVC obtained excellent performance in Germline variant calling in comparison to other widespread used software and pipelines.

**Figure 2.**
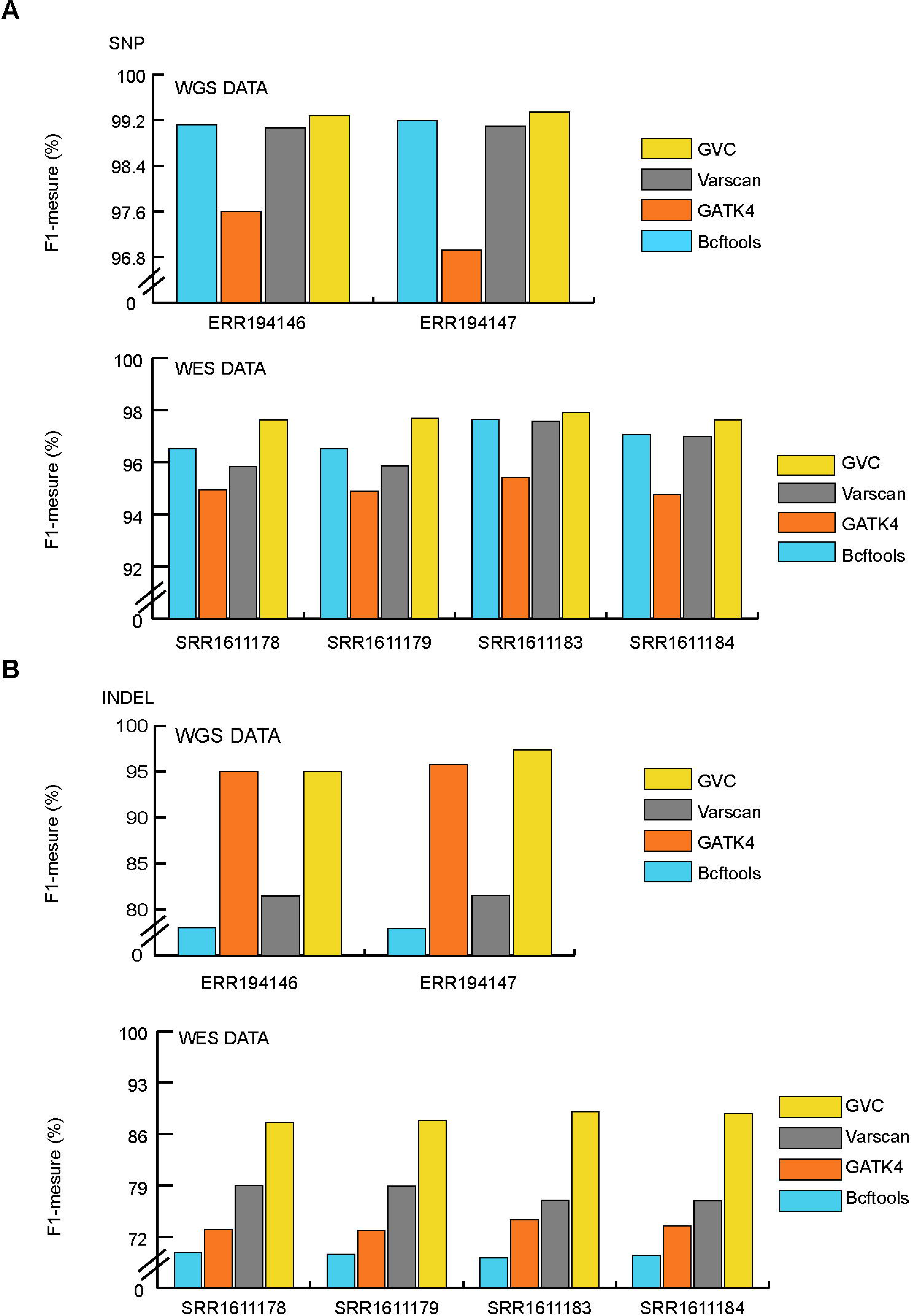
Germline variant calling performances of Bcftools, GATK4, Varscan and GVC on NA12878 and NA12877. (A) Comparison of the performances between GVC and commonly used SNP calling softwares on WGS and WES data. ERR194146 and ERR194147 are the WGS data of NA12877 and NA12878, respectively. SRR1611178, SRR1611179, SRR1611183 and SRR1611184 are all WES data of NA12878. (B) Comparison of the performances between GVC and commonly used INDELs calling softwares on WGS and WES data. ERR194146 and ERR194147 are the WGS data of NA12877 and NA12878, respectively. SRR1611178, SRR1611179, SRR1611183 and SRR1611184 are all WES data of NA12878.

### Accuracy of GVC on Somatic variant calling

We downloaded forty WGS breast cancer datasets with matched normal-tumor paired samples from the PCAWG (Pan-Cancer Analysis of Whole Genome) project on AWS (Amazon Web Services) for GVC WGS Somatic variant training. Ten-fold cross validation strategy was applied in training process (see method). We evaluated the accuracy of GVC’s SNVs and INDELs calls against the consensus dataset declared by ICGC (see method). GVC obtained mean F1-measure of 90.79% for WGS Somatic SNVs. This detection rate exceeded those of MUSE tools as well as pipelines from Sanger institution, Broad institution and German Cancer Research Center (DKZF) (SNVs: ≤ 88.12%) (Figure 3A and Supplementary Table 5). GVC achieved mean F1-measure of 76.88% for WGS Somatic INDELs. This detection rate exceeded those of pipelines from Sanger, Broad and DKZF (INDELs: ≤75.66%) (Figure 3B and Supplementary Table 6). In order to verify the robustness of the WGS Somatic model, we randomly picked 80 Pan-Cancer datasets from 8 cancer types (10 dataset for each cancer type) for testing. Results showed that the SNVs average performance is over 88% and INDLE average performance is over 70%. GVC also achieved excellent performances on Pan-Cancer WGS datasets (Supplementary Fig. 2). Taken together, GVC exhibits superior performance in WGS Somatic variants calling than other widely used software and pipelines.

**Figure 3.**
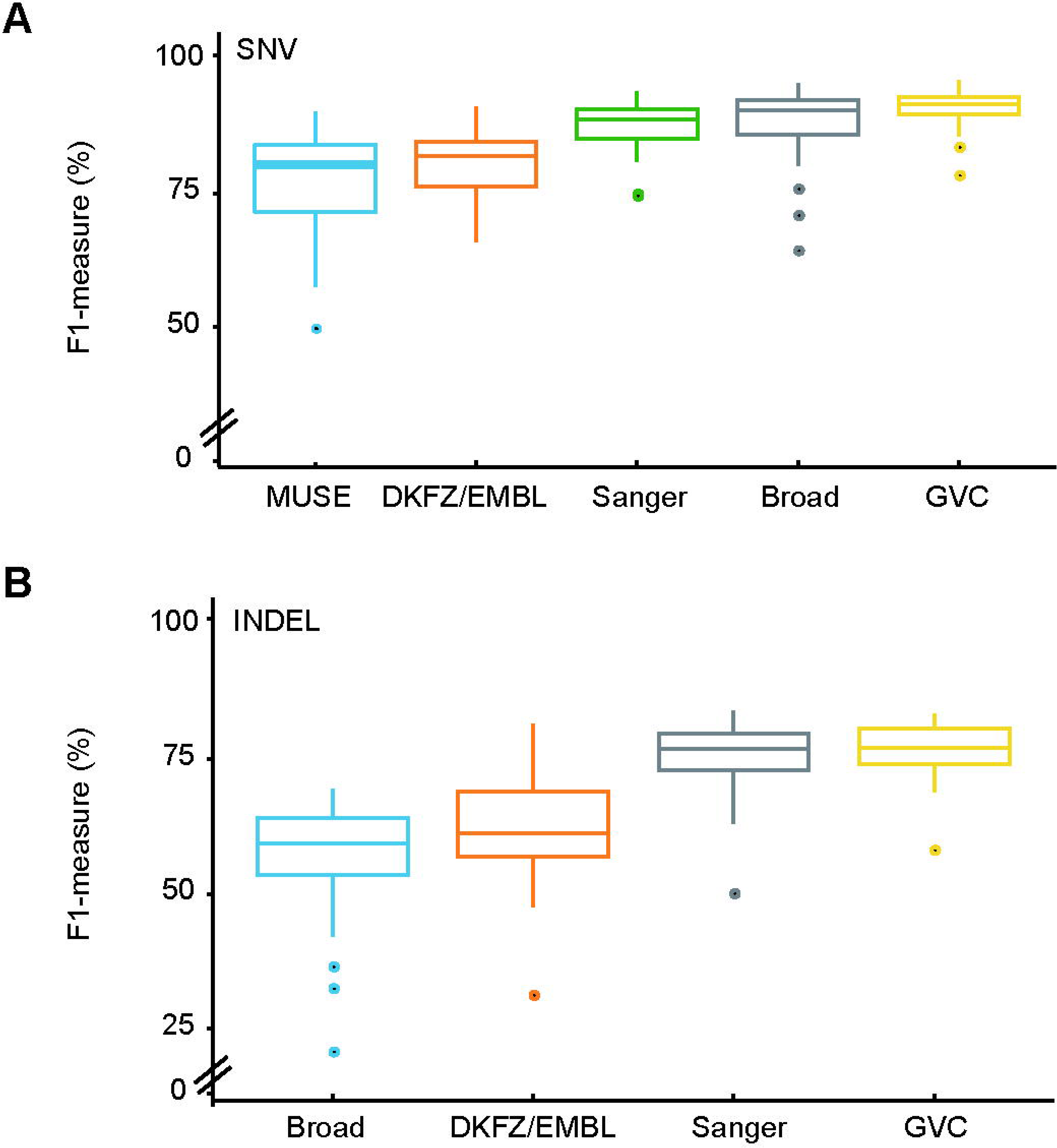
WGS Somatic variant calling performances of GVC and other softwares and pipelines on 40 WGS breast cancer dataset from ICGC. (A) Comparison of the performances between GVC and widely used SNVs calling softwares (MUSE) and pipelines from Sanger, DKFZ/EMBL and Broad institutions. (B) Comparison of the performances between GVC and widely used INDELs calling pipelines from Sanger, DKFZ/EMBL and Broad institutions.

XGBoost (eXtreme Gradient Boosting) algorithm was used for WES SNVs and INDELs model training (see method). We obtained around 7,000 WES pan-cancer datasets with matched normal-tumor pairs from TCGA (The Cancer Genome Atlas) project on GDC data portal for GVC WES Somatic variant training.

Total 1,000 training data from 11 cancer types and 846 validation data from 16 cancer types were used for GVC SNVs model training and validation, respectively (Supplementary Table 7). We evaluated the performance of GVC’s SNVs calls against the consensus dataset declared by TCGA MC3 (see method). SNVs results showed that GVC outperformed all the other widely used softwares. The mean F1-measure of 846 validation datasets was 86.3% in GVC, 65.37% in Varscan, 85.61% in MUTECT, 85.3% in MUSE, 72.14% in RADIA and 65.11% in SomaticSniper (Figure 4A and Supplementary Table 8). GVC showed excellent performance of WES SNVs calling on TCGA Pan-Cancer datasets with mean F1-measure above 86.53% (Supplementary Fig. 3A).

**Figure 4.**
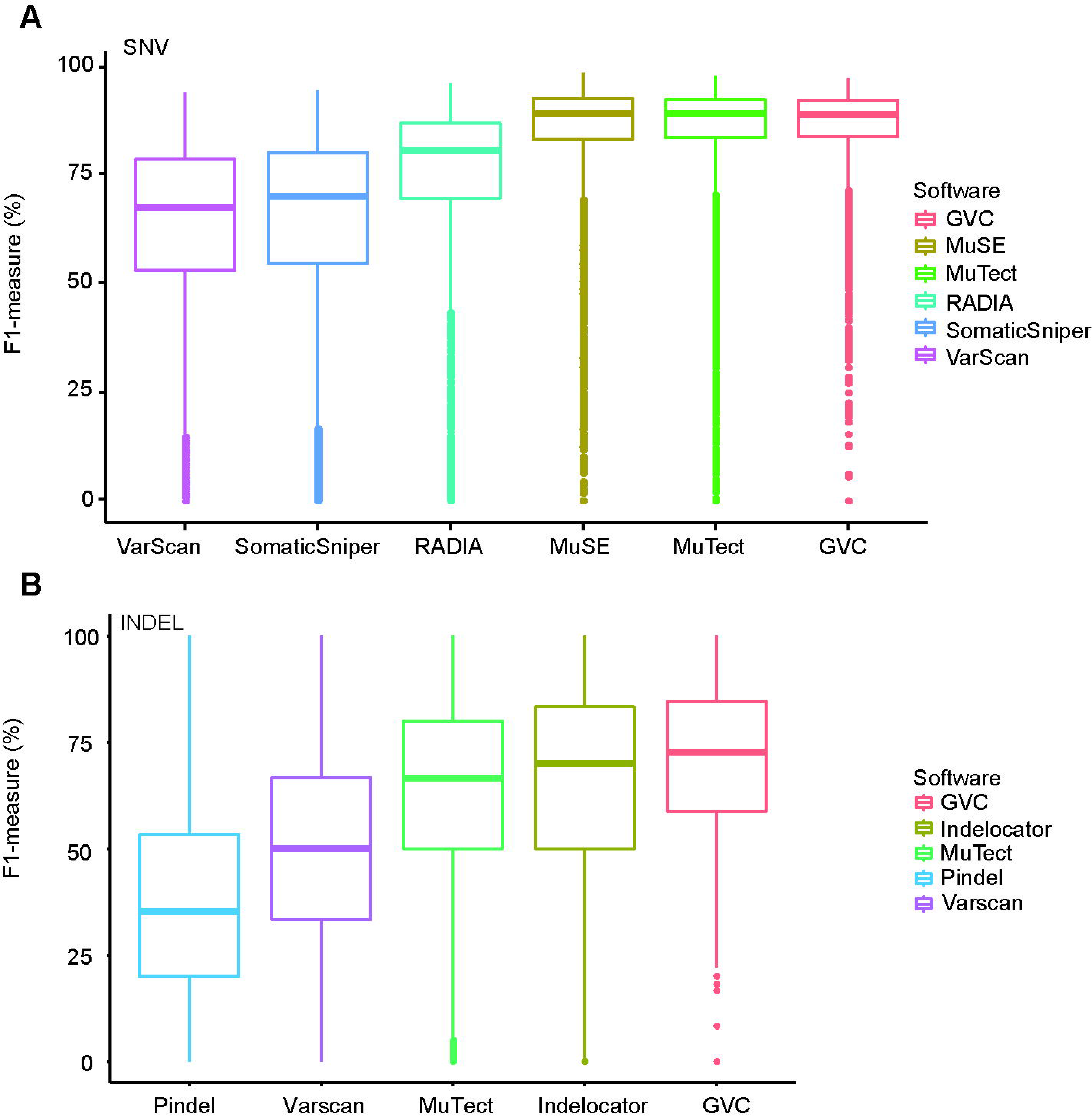
WES Somatic variant calling performances of GVC and publicly used softwares on TCGA datasets. (A) WES Somatic SNVs calling performances of GVC and publicly used softwares on TCGA datasets. (B) WES Somatic INDELs calling performances of GVC and publicly used softwares on TCGA datasets.

Total 1,701 training data from 10 cancer types and 649 validation data from 14 cancer types were used for GVC INDELs model training and validation, respectively (Supplementary Table 7). We evaluated the performance of GVC’s INDELs calls against the high confidential ANS datasets we defined (see method). INDELs calling of GVC also achieved appealing performance. Results showed that GVC achieved mean F1-measure of 68.36% for WES Somatic INDELs (Figure 4B and Supplementary Table 9). This detection rate exceeded those of Indelocator, Pindel, Varscan and Mutect tools (INDELs: ≤ 64.61%). GVC showed excellent performance of WES INDELs calling on TCGA pan cancer datasets with mean F1-measure above 73.22% (Supplementary Fig. 3B). Taken together, GVC showed superior performance in WES Somatic variants calling than other widely used softwares.

TMB (tumor mutation burden), defined as the number of somatic, coding, nonsynonymous mutations per megabase of genome examined (mutations/Mb coding region), is an evolving biomarker that may be helpful in selecting patients for immunotherapy. Accurate mutation calling is a pivotal prerequisite for TMB evaluation. Aforementioned results showed that GVC had superior performance in WES Somatic variant calling than other publicly used softwares. We further compared the total mutation load (ML) called by GVC with ML called by other MC3 softwares. It has been shown that GVC had the minimum ML deviation from the consensus MC3 dataset when comparing with other publicly used softwares (Supplementary Fig. 4). Basing on the accurate GVC algorithm, we released the TMB results of TCGA datasets on GVC Web Tool (http://gvc.0cancer.cn).

### Run time comparison

Variant detection is a time-consuming process. We compared the running time of GVC with other three widely used softwares VARSCAN, Bcftools and GATK in Germline mutation calling. WGS (average depth: 50X) and WES (average depth: 90X) datasets of NA12877 and NA12878 were used for running time testing, and the main computing resources are 20 cores and 384 memory. The results of GVC in WGS datasets showed 32-fold, 27-fold and 128-fold increase in processing speed in comparison to VARSCAN, Bcftools and GATK, respectively (Figure 5A WGS). The results of GVC in WES datasets exhibit 36-fold, 42-fold and 54-fold increase in processing speed in comparison to VARSCAN, Bcftools and GATK, respectively (Figure 5A WES). Ten WES datasets were randomly used (average depth around 100X) for running time testing in Somatic mutation calling. One thread was used to run the software and the main computing resources are 40 cores and 384G memory. The results of GVC in WES datasets showed 9.6-fold and 5.3-fold increase in processing speed in comparison to MUTECT and VARSCAN, respectively (Figure 5B). Taken together, ultra-fast processing speed and superior performance of GVC on both Germline and Somatic variant calling pave the way to its extensive application in both academic and clinical scenario. Aforementioned GVC Web Tool (http://gvc.0cancer.cn) was developed for users who can freely experience GVC variant calling performance and the compassion of TMB between user’s cohort and TCGA cohort.

**Figure 5.**
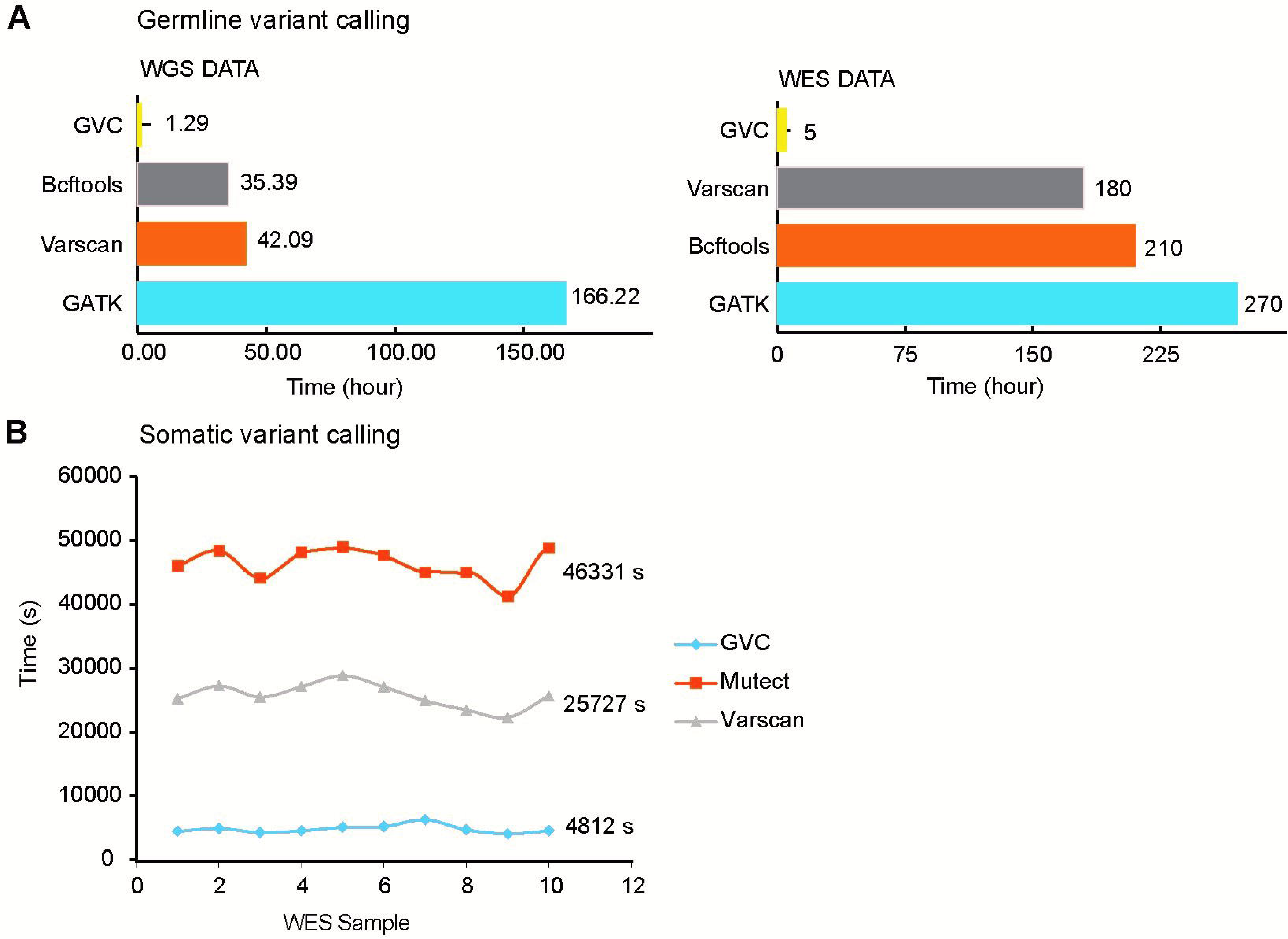
Run time comparisons between GVC and widely used softwares. (A) Comparison of the run time of Germline variant calling between GVC and widely used softwares. (B) Comparison of the run time of Somatic variant calling between GVC and widely used softwares.

## CONCLUSION

Rapidly developed NGS technology has made data analyzing a big challenge for both researchers and clinicians. How to identify mutations efficiently and accurately is still a big problem. Here we exploited GVC, combining Feature Space construction with XGBoost machine learning method, to obtain a data-driven training model for Germline and Somatic variant calling with ultra-fast speed and high accuracy. Basing on the accurate GVC algorithm, we released tumor mutation burden (TMB) results of TCGA datasets on GVC Web Tool http://gvc.0cancer.cn.

## METHODS

### Datasets for Germline mutation

We used the NA12877 and NA12878 datasets for Germline variant training and testing. They are the most studied genome came from Coryell CEPH/UTAH 1463 family. We downloaded the WGS data from Platinum Generals Project (Https://www.illumina.com/platinumgenomes.html), where the sequencing platform is Illumina Hiseq 2000 and the depth is 50X. In the paper, we chose four WES samples, including SRR1611178, SRR1611179, SRR1611183 and SRR1611184, among them SRR1611178 and SRR1611179 came from Illumina Hiseq 2000, while others came from Illumina Hiseq 2500.

During Germline model training, we used only the NA12878 data set. We used a high confidential set of NA12878 which published by GIAB (Genome in a bottle consortium) to mark the train set. We used the rtg-tools which can process complex VCF format files, detect the variation location, assess mutants Genome type (heterozygous or homozygous), and evaluate the performance within a highly confidential intervals.

### Datasets for Somatic mutation

The PCAWG (Pan-cancer Analysis of Whole Genomes) projects on AWS include more than 2,400 genomes from 1100 donors, among which have 44 breast normal-tumor paired WGS samples. We abandoned four sets of data with low consistency in INDELs consensus, and finally used 40 sets of data for model training and testing.

We selected 40 sets of ICGC breast data, using the consensus published by ICGC. ICGC consensus sets are built by variant calling pipelines from four academic organizations which respectively are the German Cancer Research Center (DKFZ), the European Molecular Biology Laboratory (EMBL) in Heidelberg, the Wellcome Trust Sanger Institute, and the Broad Institute. In order to verify the robustness of the WGS Somatic model, we pick eight cancer projects form PCAWG, each randomly selects 10 datasets for performance evaluation.

ICGC annotates consensus results for different situations, such as PASS, LOWSUPPORT (Not called by enough callers in ensemble), OXOGFAIL (Failed OXOG oxidative artifact filter), bSeq (Sequencing Bias), bPcr (PCR Bias), GERM1000G (1000Genome variant with insufficient somatic evidence), GERMOVLP (Overlaps germline Haplotype call), NORMALPANEL (Presence in Panel of Normals), REMAPFAIL (Variant no longer seen under remapping), SEXF (Likely artifact or call in PAR region: Y-chromosome variant in female donor). We chose the PASS filter label as the SNP “ground truth” set. Unlike the consensus result of SNVs which established by four academic pipeline, ICGC’s INDELs consensus result comes from three variant calling pipeline, including Sanger, Broad and DKFZ. “LOWSUPPORT” variants called by single software would be credible through IGV. So the INDELs “ground truth” set including (1) marked as PASS point set; (2) Single software supported and tumor Alt count >=4, normal alt count <=0. We use the “ground truth” set to mark training data and evaluate predictive results.

Around 7,000 pan-cancer datasets with matched normal-tumor pairs from TCGA were employed for WES Somatic variant training with XGBoost (eXtreme Gradient Boosting) algorithm. Total 1,000 training data from 11 cancer types and 846 validation data from 16 cancer types were used for GVC SNVs model training and validation, respectively, and total 1,701 training data from 10 cancer types and 649 validation data from 14 cancer types were used for GVC INDELs model training and validation, respectively.

### Measure method

The following 4 measures were used to estimate classifier performances:

1. Sensitivity (or Recall or true positive rate) measures the proportion of the known Somatic variants those are correctly predicted as those and is defined as TP/ (TP + FN), where TP is true positive and FN is false negative.
2. Precision is a fraction of the correctly called Somatic mutations to all variants that are labeled as Somatic by the classifier and is defined as TP/ (TP + FP), where FP is false positive.
3. F1-measure is the harmonic mean of precision and recall: 2 × (Precision × Recall)/ (Precision + Recall).
4. Area under ROC curve (AUC) denotes the probability that a classifier assigns a higher score to the positive instance than a randomly chosen negative sample. It measures the general ability of the classifier to separate the positive and negative classes. The best performing classifier for each cancer dataset was selected based on AUC and F1-measure.

### Feature construction

GVC extracts feature information from BAM file to assemble feature vector space, which includes: (1) the quality statistics information, such as the mapping quality and base quality; (2) some number statistics, such as mismatch number, read depth, read number with different start point, read strand, mutant frequency; (3) the distance information, such as the distance from variant point to read end; (4) Database Library, such as dbSNP, COSMIC and 1000G frequency; (5) Features are constructed according to other SNP software such as GATK and Samtools, such as Consensus base, SNP quality, root mean square mapping quality and Phred-scaled p-value and Somatic score; (6) Other statistic, such as Fisher test.

### Training method

Large amount of training set leads to large memory consumption and long training time. Meanwhile, too many negative loci will affect the accuracy of the model. So we did the pretreatment: In Germline training, satisfying the condition that read_depth >= 10, Mutant read number >= 2, baseQ >= 20 and mapQ >= 20 are chose as the candidate set. In WGS Somatic training, the training set must satisfy the condition that read_depth >= 10, Mutant read number >= 2, normal frequency <= 10%, baseQ >= 20 and mapQ >= 20. In the training step, we adopted XGBoost method, a machine learning algorithm based on gradient big regression tree. In order to make full use of the data, we adopted the k-fold cross validation training strategy.

And for WES Somatic training, XGBoost used the maximum threshold of trees 5,000, and the depth of the tree 3. During model construction, 80% and 20% of the datasets were randomly utilized for model training and inner validation, respectively. And the training model was finally tested for its performance in separate testing datasets.

## SUPPLEMENTARY DATA

Supplementary data are available at NSR online.

## Acknowledgments

We gratefully acknowledge contributions from the Platinum Genomes Project, PCAWG, GIAB, ICGC and TCGA and the specimen donors.

## Funding

This work was supported by no funding.

## Author contributions

Hao L., Zhang Z., Cao L., and Ling S. designed the core algorithms; Yin L., and Zhang Z. performed Feature construction; Liu C., and Zhang Z. selected the training model; Zhang Z. conceived the training project and performed the analysis of ICGC datasets; Hao L., Chen M., and Zhang Z. conceived the training project and performed the analysis of TCGA datasets; Li C., and Xu Z., and Ling S. supervised the project designing and data analysis processes; Zhang Z., Chen M., Xu Z., and Ling S. wrote the manuscript; All authors discussed the results and revised the manuscript.

### Competing financial interests

The authors declare no competing financial interests.

## SUPPLEMENTARY FIGURE LEGENDS

**Supplementary Figure 1.**
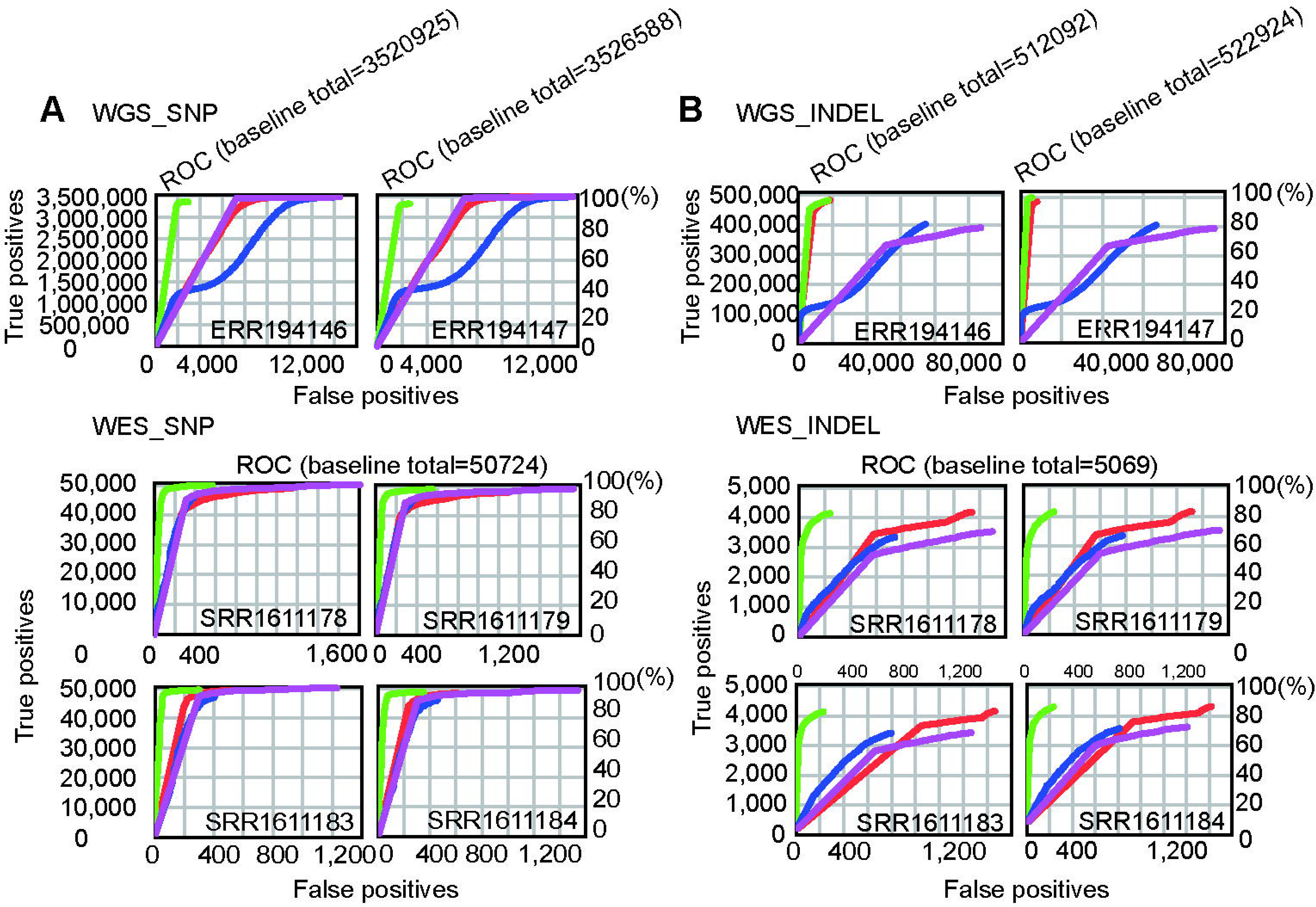
ROC curves for method comparison. Data used to construct these ROC curves were obtained from Bcftools (purple), GAKT4 (red), Varscan (blue) and GVC (green). ROC curve is built by rtg-tools.

**Supplementary Figure 2.**
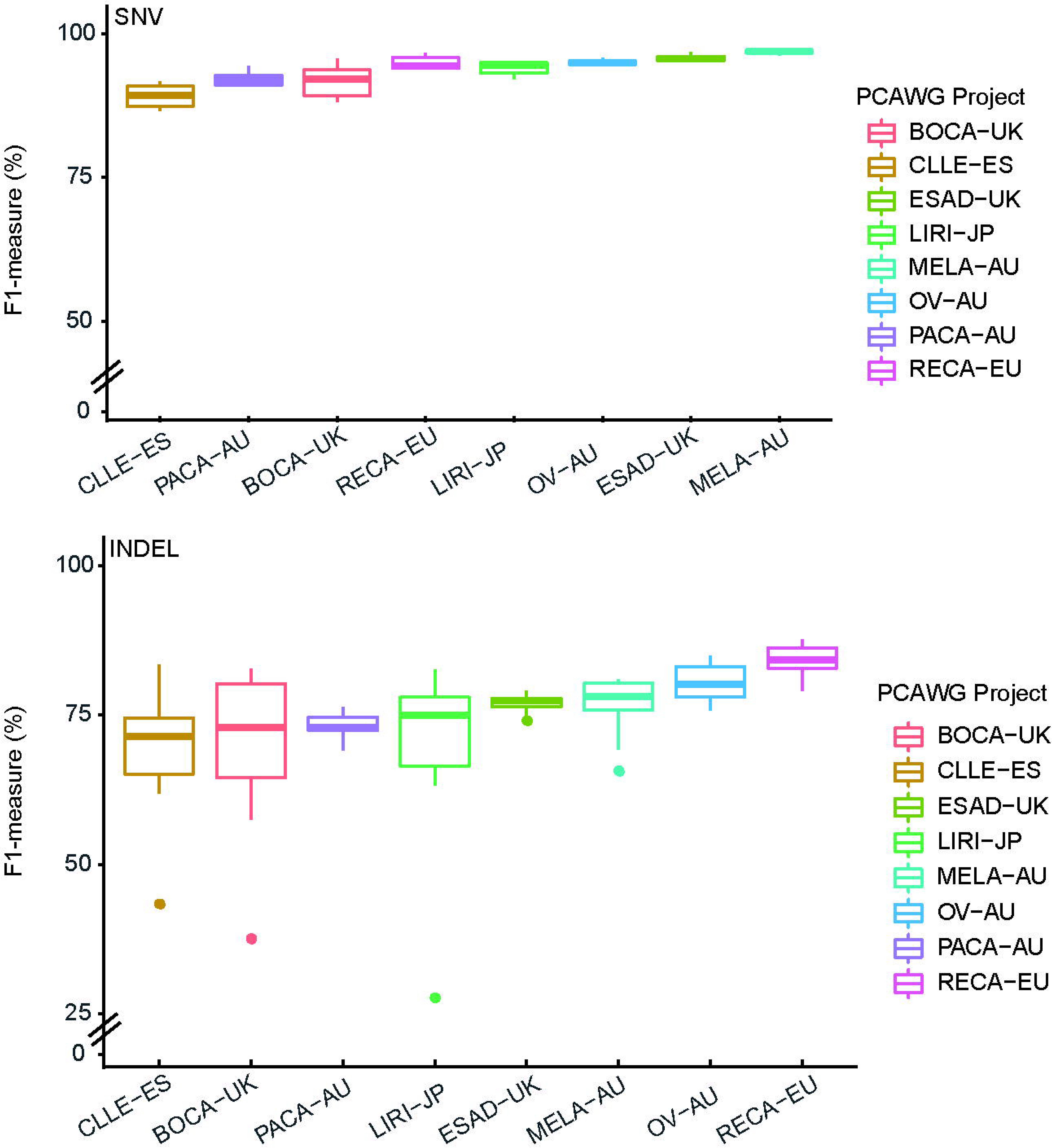
Robustness of GVC WGS Somatic model.. Using breast cancer training model to predict Pan-Cancer datasets (80 datasets from 8 cancer type) from PCAWG Project.

**Supplementary Figure 3.**
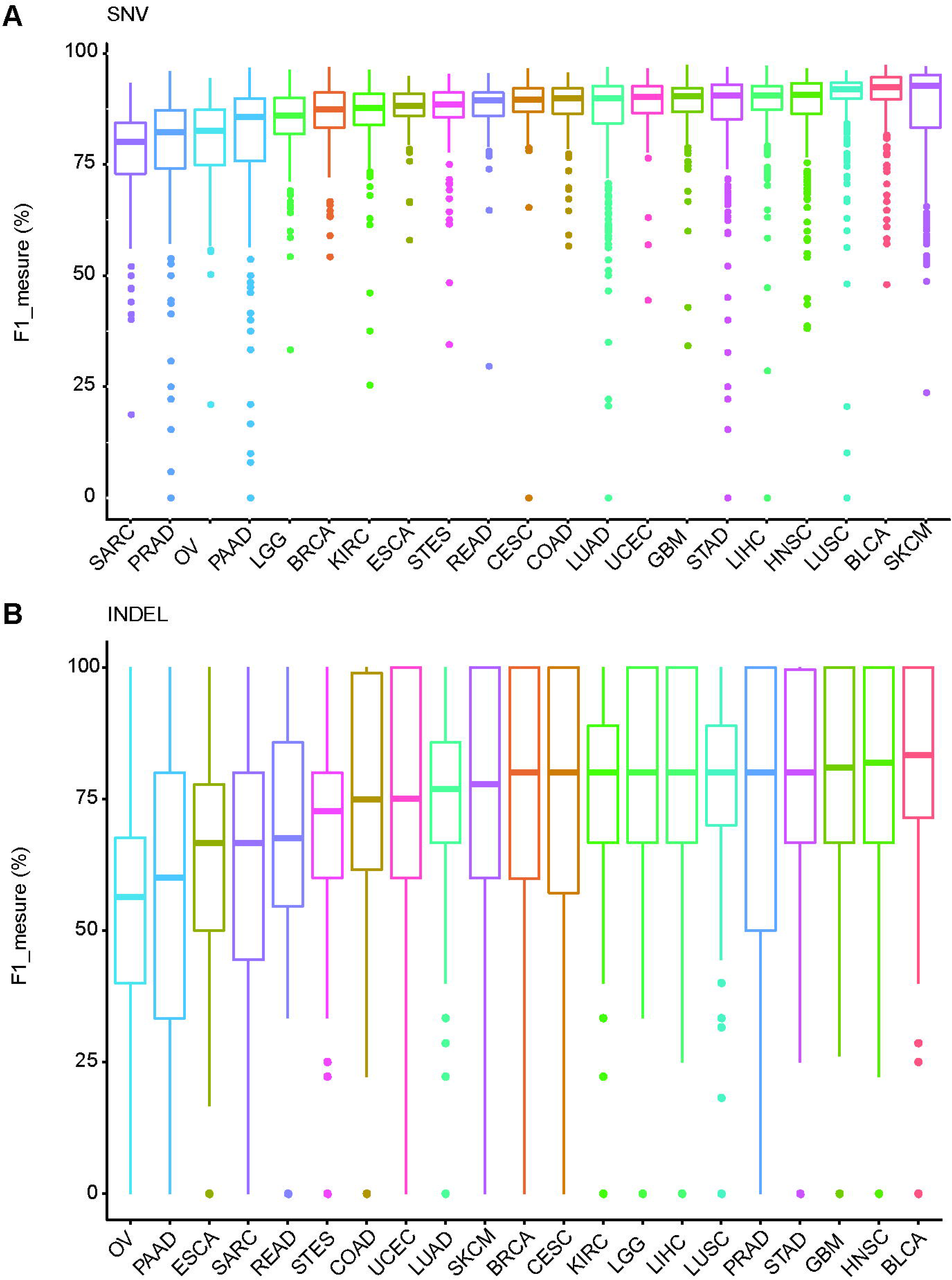
The performance of GVC on TCGA pan cancer datasets. (A) The performance of GVC of WES SNVs calling on TCGA pan cancer datasets. (B)The performance of GVC of WES INDELs calling on TCGA pan cancer datasets.

**Supplementary Figure 4.**
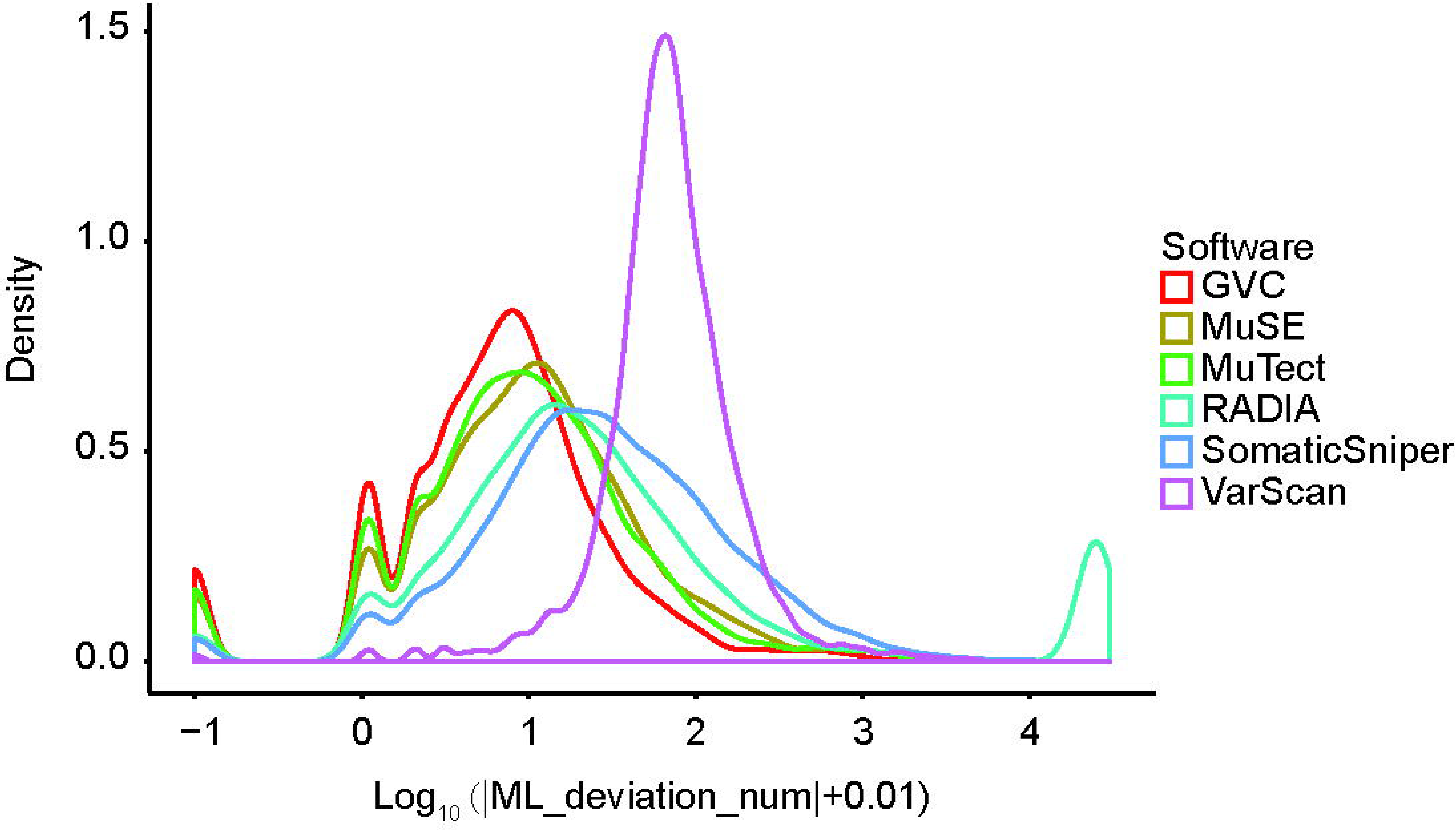
Mutation load (ML) deviation of GVC and other publicly used softwares from the consensus MC3 datasets. ML deviation numbers between ML of softwares and ML of MC3 were calculated. And Log_10_ (|ML_deviation_num| + 0.01) was used for density plot.

**Table S4:**
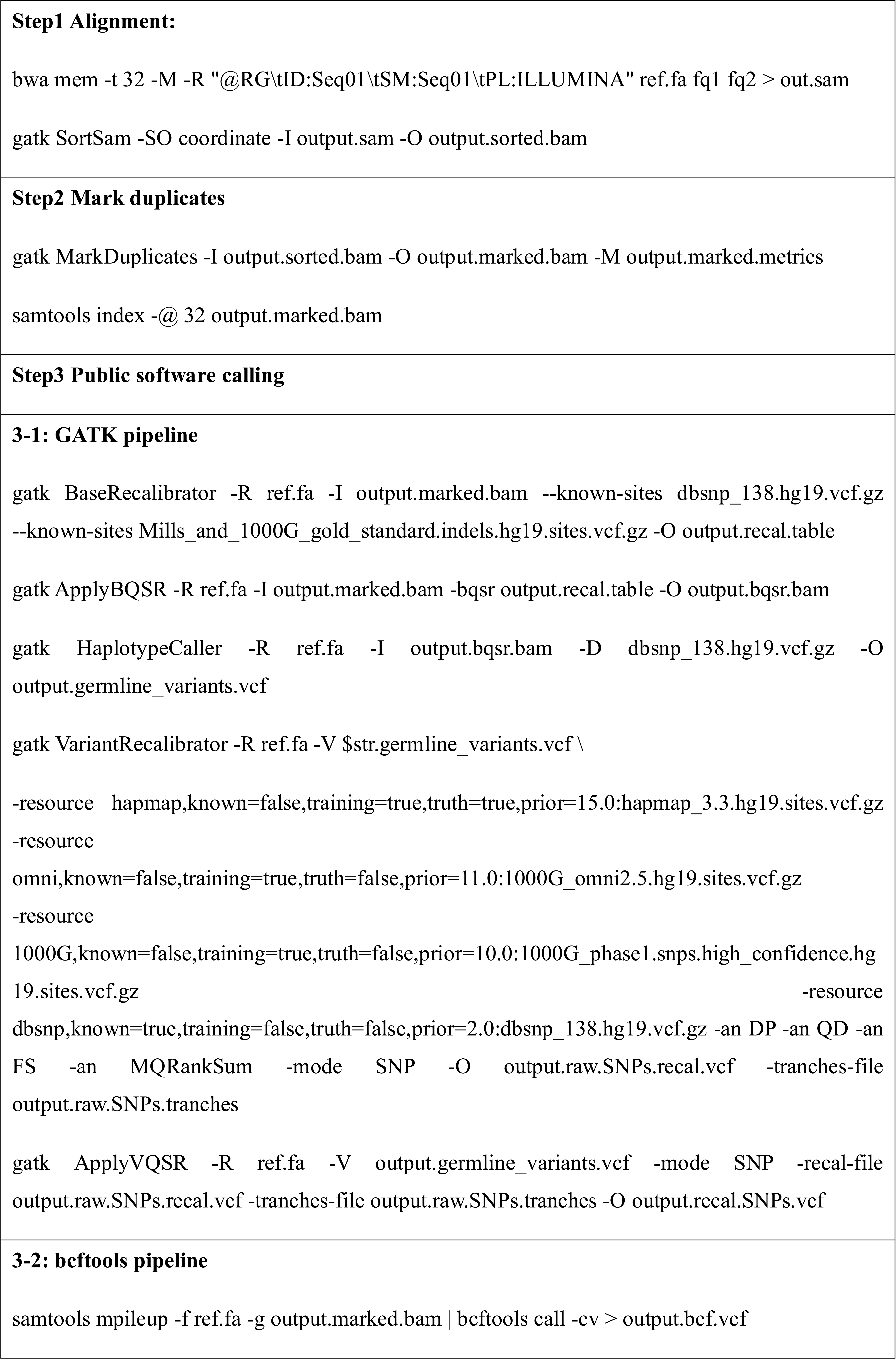
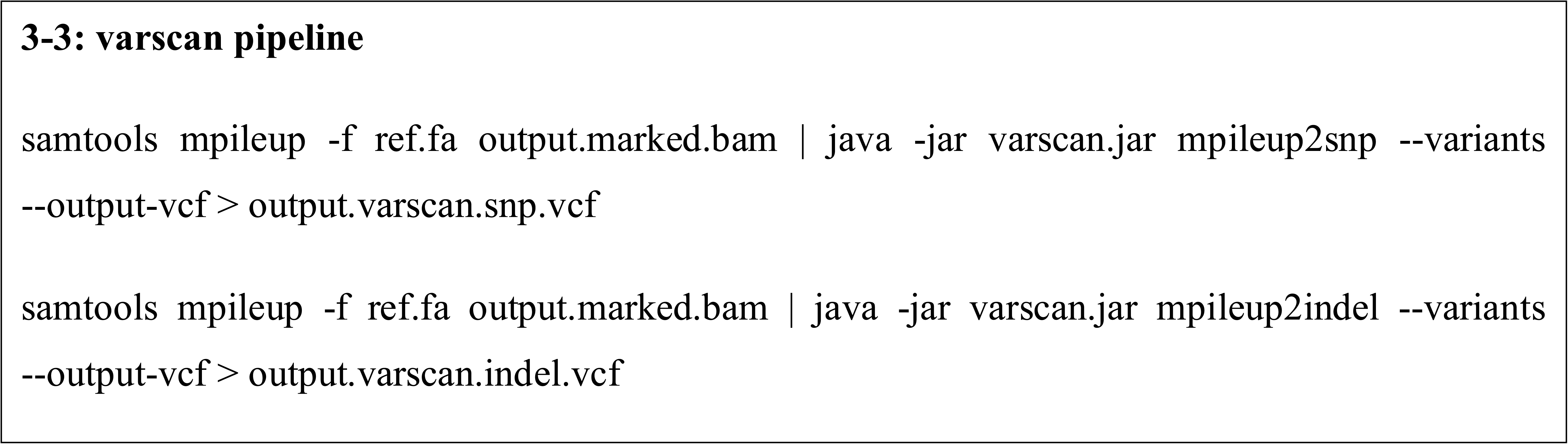
Scripts used in Germline variant calling of GATK, Bcftools and Varscan.

